# UNC-45A is preferentially expressed in epithelial cells and binds to and co-localizes with interphase MTs

**DOI:** 10.1101/615187

**Authors:** Juri Habicht, Ashley Mooneyham, Mihir Shetty, Xiaonan Zhang, Vijayalakshmi Shridhar, Boris Winterhoff, Ying Zhang, Jason Cepela, Timothy Starr, Emil Lou, Martina Bazzaro

## Abstract

UNC-45A is a ubiquitously expressed protein highly conserved throughout evolution. Most of what we currently know about UNC-45A pertains to its role as a regulator of the actomyosin system. However, emerging studies from both our and other laboratories support a role of UNC-45A outside of actomyosin regulation. This includes studies showing that UNC-45A: regulates gene transcription, co-localizes and biochemically co-fractionates with gamma tubulin and regulates centrosomal positioning, is found in the same subcellular fractions where MT-associated proteins are, and is a mitotic spindle-associated protein with MT destabilizing activity in absence of the actomyosin system.

Here, we extended our previous findings and show that UNC45A is variably expressed across a spectrum of cell lines with the highest level being found in HeLa cells and in ovarian cancer cells inherently paclitaxel-resistant. Furthermore, we show that UNC-45A is preferentially expressed in epithelial cells, localizes to mitotic spindles in clinical tumor specimens of cancer and co-localizes and co-fractionates with MTs in interphase cells independent of actin or myosin.

In sum, we report alteration of UNC45A localization in the setting of chemotherapeutic treatment of cells with paclitaxel, and localization of UNC45A to MTs both *in vitro* and *in vivo*. These findings will be important to ongoing and future studies in the field that further identify the important role of UNC45A in cancer and other cellular processes.

## Introduction

The uncoordinated protein 45 (UNC-45) is ubiquitously expressed and highly conserved throughout evolution[1, 2]. Vertebrates have two UNC45 isoforms: UNC45B which is expressed only in muscle cells, and UNC45A which is expressed in all cells. While the conservation of UNC-45 suggests its critical importance in cell and developmental biology, as well as pathophysiologic processes such as cancer, UNC-45 functions are still largely unknown. Structurally, UNC-45 can be divided in four domains: an N-terminal domain, which contains three tetratricopeptide repeat (TPR) sequences and has been shown to bind to Hsp90; a central domain of largely unknown function; a neck domain, which has been recently proposed to be required for UNC-45 oligomerization[3]; and a C-terminal, UCS domain which is the most well studied and that is critical for interaction with the motor domain of myosin [2, 4]. Identifying spatial localization of UNC45A in cells and also layout in the context of human tissue has potential strong implications for pathophysiology of cancer cell development and development of chemotherapy resistance of dividing cells. Thus, we hypothesized that UNC45A localizes to the microtubule complex and that validated anti-UNC45A antibodies could effectively be utilized to stain regions of UNC45A intensity for microscopic visual examination. Here, we report our findings that UNC45A is preferentially expressed in cells of epithelial origin, and localizes to the MT complex *in situ* in vivo in human tumor tissues as well as *in vitro*.

## Results

### Validation of anti-UNC45A antibody by Western blot analysis. Full membranes

In our work to date, we have used several anti-UNC45A antibodies, from different source companies. We identified the need to streamline use of anti-UNC45A antibodies testing for future studies, and thus sought to provide a biochemical validation of two of the most commonly used commercially available UNC-45A antibodies: the rabbit polyclonal anti-UNC-45A from Proteintech, which we have previously used to show that UNC-45A is a mitotic spindle-associated protein [5], and the mouse polyclonal anti-UNC-45A from Abnova, which was used to show that UNC-45A is required for stress fibers formation [6].

As shown in Figure 1A, full membrane Western blot, revealed that the Proteintech antibody detects a single band corresponding to UNC-45A in lysates of HeLa (human cervical cancer) and RFL-6 (rat lung fibroblast) cell lines. As show in Figure 1B, full membrane Western blot using the Abnova antibody also detected a band corresponding to UNC-45A in lysates of RFL-6 and HeLa cell lines, with a few other additional bands being visible, especially when using long exposure times.

**Figure 1.**
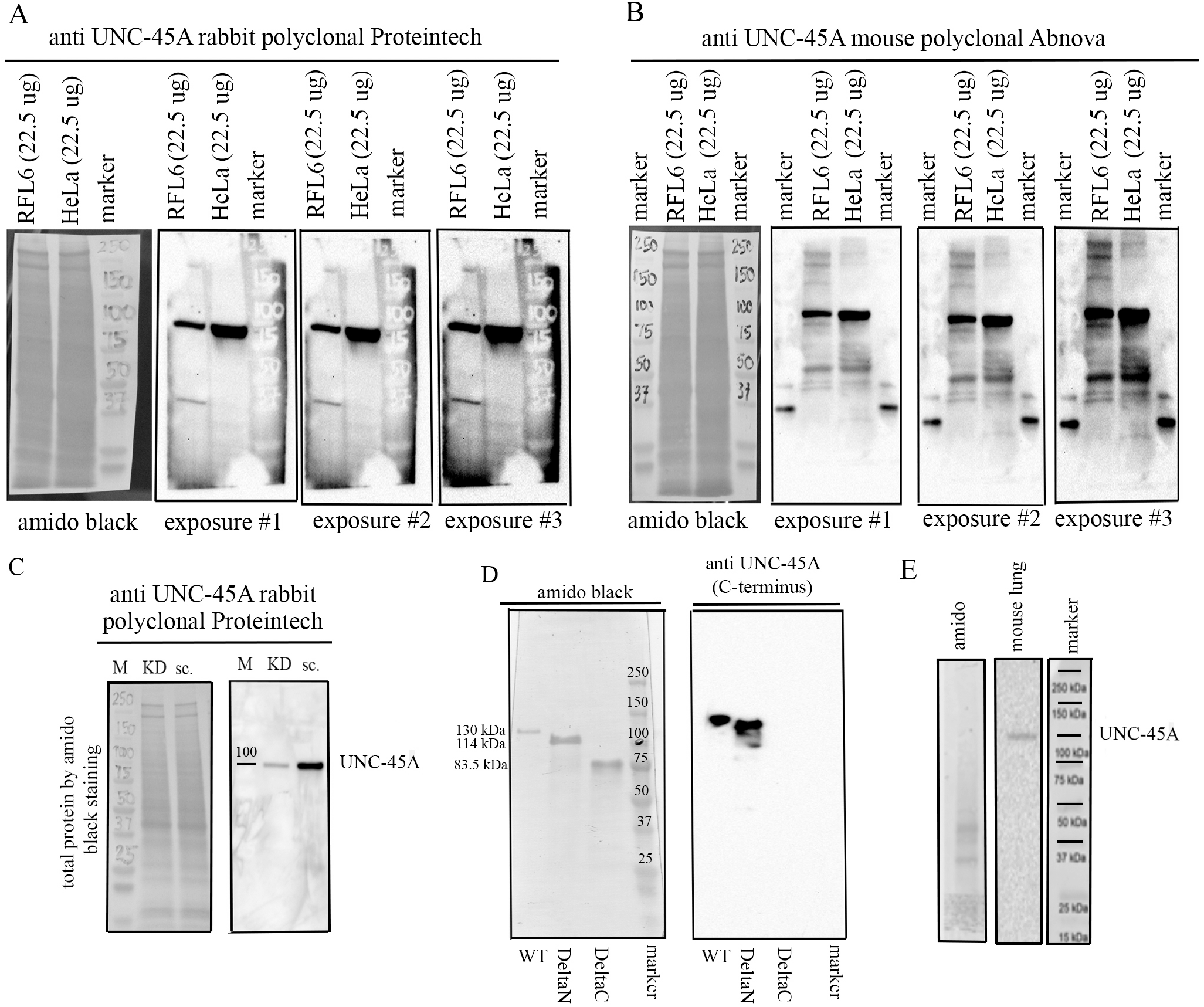
Comparison between the anti-UNC45A rabbit polyclonal and the anti-UNC-45A mouse polyclonal antibodies by Western blot. Full membranes. A. Full membrane probed with the anti UNC-45A rabbit polyclonal antibody from Proteintech shows one band at the expected molecular weight of UNC-45A (103 kDa) in lysates of RFL6 and HeLa cell lines. Amido black for equal protein loading and three different exposures are shown. Original, non contrasted images are shown. B. Full membrane probed with the anti UNC-45A mouse polyclonal antibody from Abnova shows one band at the expected molecular weight of UNC-45A (103 kDa) and few additional bands in lysates of both RFL6 and HeLa cell lines. Amido black for loading control and three different exposures (same used for the Proteintech antibody) are shown. Original non contrasted images are shown. For these experiments we overloaded the gels with 22.5 μg of total protein to ensure detection of all bands. Same samples were used for both Western blots. C. Full membranes probed with the anti UNC-45A rabbit polyclonal antibody from Proteintech in lysates (12.5 μgs) of HeLa cells 48h after transduction with either shRNA-scramble (sc.) or shRNA targeting UNC-45A (KD). Amido black was used as a loading control. Original non contrasted images are shown. D. Recombinant full length (WT, aa 1-944) UNC-45A-GFP (MW 130 kDa), deltaN (aa 125-944) UNC-45A-GFP (MW=114 kDa), and deltaC (aa 1-514) UNC-45A-GFP (83.5 kDa) were separated on a 4-15% SDS gel, transferred onto PVDF membrane and stained with amido black (*left*) or blotted against the anti UNC-45A rabbit polyclonal antibody from Proteintech. E. Full membrane probed with the anti UNC-45A rabbit polyclonal antibody from Proteintech in lysates (7.5 μgs) of adult mouse lung. Amido black is shown. To our knowledge, this is the first report showing full membranes Western blot probed with anti-UNC-45A antibodies.

We then proceeded to confirm that the single band observed with the Proteintech antibody was indeed corresponding to UNC-45A. To this end, UNC-45A was knocked down in HeLa cells via lentiviral mediated delivery of either shRNA scramble or shRNA-UNC-45A and the specificity of the rabbit polyclonal antibody was confirmed via Western blot analysis. As shown in Figure 1C, UNC-45A knockdown leads to an approximate 90% reduction in the band corresponding to UNC-45A using the rabbit polyclonal antibody.

Because the Proteintech antibody was raised against the C-terminus of human UNC-45A (as indicated by the manufacturer), we also tested its specificity on recombinant WT, deltaN and deltaC UNC-45A. As shown in Figure 1D, the antibody failed to recognize the recombinant protein lacking the C-terminal domain. Additionally, we tested this antibody in tissue lysates of mouse lung tissue as recommended by the antibody manufacturer, to confirm that this specificity of the antibody was to the C-terminus portion of UNC45A. As shown in Figure 1E, the antibody recognized a band corresponding to UNC-45A.

In sum, we found a high level of specificity of two commercially available antibodies to UNC45A. Both antibodies tested were remarkably specific and suitable for further use in experiments aimed at assessing localization of UNC45A in cells *in vitro*, *in situ*, and *in vivo*. However, because the Proteintech antibody recognizes a single band on the Western blot analysis of cellular lysates, we believe the Proteintech antibody is particularly suitable for UNC-45A in cells and tissues.

### Expression levels of UNC-45A across cell types and its co-fractionation with microtubules (MTs) in cells

Proteomic analysis of HeLa cells shows that not only is UNC-45A in the subcellular fractions where other well-known MT-associated and destabilizing proteins are found, but that it is approximately 20-fold more abundant (0.4 uM) than other MT destabilizing proteins [7]. In light of our recent report that UNC-45A is a MT destabilizing protein [5], this suggests that UNC-45A may have a dominant role in the regulation of cellular MTs and that UNC-45A levels may very across multiple cell types. Here we set forth to determine endogenous UNC-45A protein levels by Western blot analysis in commonly used cell lines including the cervical cancer cell line HeLa, the ovarian cancer cell line SKOV-3, the rat fibroblast RFL-6 cell line and in a panel of ovarian cancer cell lines: ES-2, TOV-21G, JHOC5, OVTOKO, RMGI and ES-2. HeLa, SKOV-3 and RFL-6 cells where chosen because they are most commonly used to study MT destabilizing proteins owing to their relatively flat and stable morphology and low MT density [7–9] This particular panel of ovarian cancer cell lines was chosen because they are all derived from the paclitaxel resistant ovarian cancer histotype clear cell carcinoma [10]. As shown in Figure 2A and B, there was variability in terms of UNC-45A expression levels in the different cell lines with HeLa expressing the highest levels. Noteworthily, higher UNC-45A expression levels were found in the OVTOKO, RMGI and ES-2 ovarian cancer cell lines which the ones characterized by the highest levels of chemoresistance to paclitaxel [10, 11]. This is consistent with our recently published report suggesting that UNC-45A could be a marker of chemoresistance [5].

**Figure 2.**
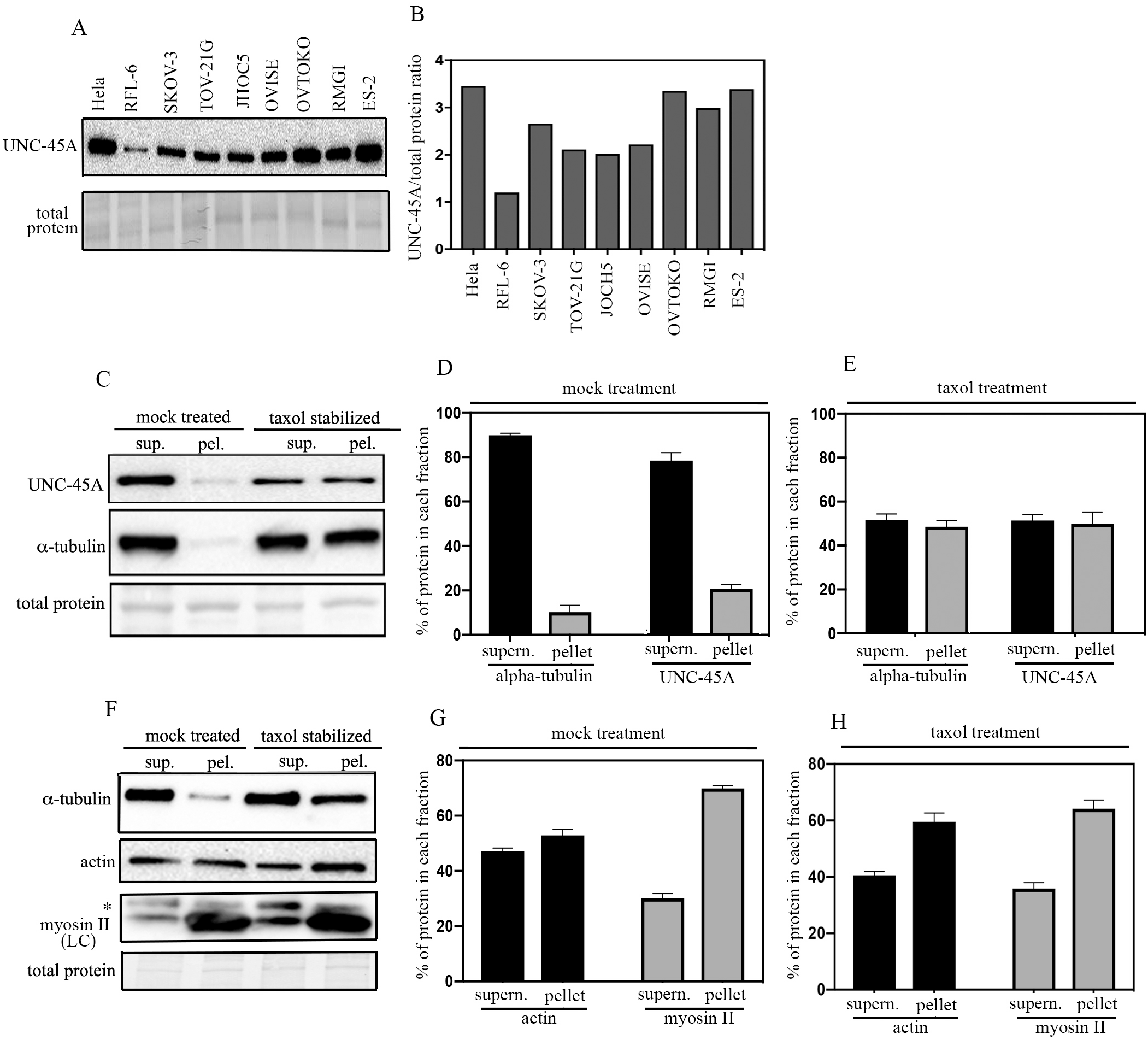
UNC-45A expression levels across cell types and its fractionation with MTs in mammalian cells. A. Western blot analysis for levels of UNC-45A in the indicated cell lines. B. Quantification of UNC-45A expression levels expressed as UNC-45A/total protein ratio. C. MTs from SKOV-3 cell lines in presence or in absence (mock) of 1µM taxol for 1 hour prior ultracentrifugation. Supernatant and pellet fractions were fractionated by SDS-PAGE and subjected to Western blot analysis using anti-UNC-45A and anti-α-tubulin, antibodies. D. Quantification of % of tubulin and UNC-45A in each fraction following mock treatment. E. Quantification of % of tubulin and UNC-45A in each fraction following taxol treatment. Overall, taxol treatment resulted in an approximately 5-fold increase in the amounts of tubulin and of UNC-45A found in the pellet versus mock. F. Actin and myosin II from SKOV-3 cell lines in presence or in absence (mock) of 1µM taxol for 1 hour prior ultracentrifugation. Supernatant and pellet fractions were fractionated by SDS-PAGE and subjected to Western blot analysis using anti-actin and anti-myosin II antibodies. Asterisk indicates an aspecific band using a monoclonal antibody against myosin II LC. G. Quantification of % of actin and myosin II in each fraction following mock treatment. H. Quantification of % of actin and myosin II in each fraction following taxol treatment. Overall, taxol treatment resulted in a slight increase (25% after adjusting for total protein loading, which is slightly higher in the bottom last lane of Figure 2F) of actin and no increase in the amount of myosin II in the pellet fractions as compared to mock treatment.

We had previously shown that endogenous UNC-45A binds to polymerized MTs and not to free tubulin in COV-362 ovarian cancer cells [5]. Here we used a standard co-sedimentation assay, which is one of the most commonly used approaches for characterizing and quantifying the ability of a protein to bind MTs [12], to determine whether endogenous UNC-45A cofractionates with MTs in living cells. In this assay, cells are treated (or not, vehicle) with taxol so to increase the fraction of MTs and microtubule-associated-proteins (MAPs) and the NP-40-resistant cytoskeletal ghosts are separated from the soluble fraction via ultracentrifugation. The fraction of each protein in supernatant and pellet fraction is then determined via Western blot analysis (Figure 2C). As shown in Figure 2D, in absence of taxol, most of the tubulin and of UNC-45A are found in the supernatant. For tubulin, this is consistent with the fact that in living cells, MTs are both dynamically unstable and quite sensitive to manipulation [13]. Taxol treatment however results in an approximatively 5-fold increase in the amounts of tubulin and of UNC-45A found in the pellet which correspond to an increase in MTs versus free tubulin (Figure 2E). We next performed the same experiment and quantified the % of actin and myosin II in each fraction with or without taxol treatment (Figure 2F). As shown in Figures 2G in absence of taxol (mock treatment), actin is present in both the supernatant and pellet in similar proportions while almost 70% of the myosin II is present in the pellet. Addition of taxol, results in a slight increase (25% after adjusting for total protein loading, which is slightly higher in the bottom last lane of Figure 2F) of actin and no increase in the amount of myosin II in the pellet fractions (Figure 2H). Taken together, this indicates that the increased UNC-45A in the pellet fraction following taxol-treatment is due to its binding to MTs and not to actomyosin.

### UNC45A is expressed in cytoplasm and mitotic spindles and is preferentially expressed in epithelial cells

We and other have previously shown that UNC-45A is a mitotic spindle-associated protein in a number of cell types including cancer cells and fibroblasts and that it has a predominantly cytoplasmic subcellular localization [5, 14–16]. We and others have also previously shown that UNC-45A is overexpressed in cancer versus benign tissue and in cancer cell lines versus normal cells [14, 17]. Here we wanted to determine the localization pattern of UNC-45A *in situ*, using the same anti-UNC-45A Proteintech antibody we had previously validated by performing IHC staining in shRNA scramble versus shRNA knockdown cells (Supplementary Figure 1[5]). As shown in Figure 3A, we found that in clinical specimens ovarian cancer, UNC-45A is present at the mitotic spindle of metaphase ovarian cancer cells *in situ*. This is consistent with our previous report of UNC-45A localization with MTs including with the mitotic spindle in multiple synchronized cell lines including HeLa, COV-362 cancer cells and NIH3T3 fibroblasts [5]. We also found that in interphase cells *in situ* UNC-45A has mostly cytoplasmic localization, and is predominantly expressed in epithelial ovarian cancer cells versus stroma which is mostly composed of fibroblasts (Figure 3B). Not only is this consistent with UNC-45A overexpression in cancer versus normal cells, but also with the fact that the cell line RFL-6, which is of fibroblastic origin, expressed the lowest levels of UNC-45A (Figure 2A and B). Next, we analyzed a single cell RNA sequencing (scRNAseq) dataset consisting of ∼90,000 single cells from 45 ovarian cancer tissue samples, to determine UNC-45A RNA expression levels in cell types present within the tumor environment. Cells were clustered based on global RNA expression using a graph-based clustering method [18] and assigned a cell type based on marker genes (Figure 3C left panel). Based on analyses of these data, UNC45A is expressed at a higher level in epithelial cells compared to stromal and immune cells (Figure 3C left panel and Figure 3D). Taken together, this suggests that expression of UNC-45A is predominantly found in epithelia and cancer cells and may be associated with their higher proliferative status.

**Figure 3.**
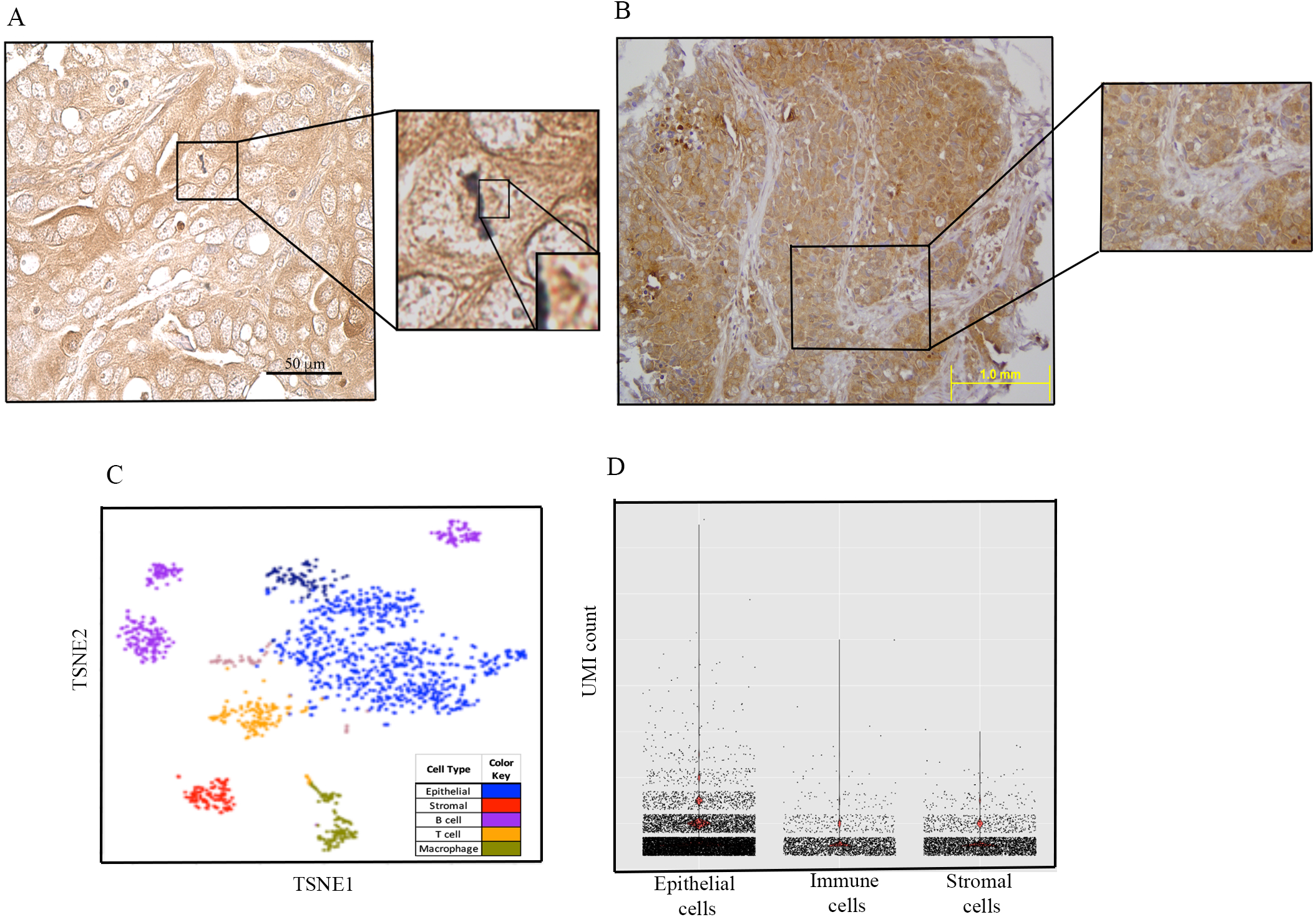
Pattern of UNC-45A expression *in situ*. A. Immunohistochemical staining of UNC-45A in clinical specimens ovarian cancer shows UNC-45A localization on mitotic spindle in a metaphase cancer cell *in situ*. B. Immunohistochemical staining of UNC-45A in clinical specimens ovarian cancer shows that UNC-45A localization is mostly cytoplasmic in cancer cells and that its levels are higher in cancer cells versus stromal cells. C. Clustering of 1,492 single cells from a representative sample (Patient #101), colored by cell type (left panel) or by UNC45A expression (right panel). D) Barplot of UNC45A expression in 90,000 epithelial, stromal and immune cells from 45 ovarian tissue samples. UNC45A expression is measured based on unique molecular identification (UMI) barcode counts.

### UNC-45A colocalizes with interphase MTs *in vivo*

We have shown that UNC-45A binds to purified MTs *in vitro* and to mitotic spindles in cells with a punctated pattern [5]. This pattern is consistent with which is consistent with what has been shown for other MT destabilizing proteins [5, 8]. Here we wanted to determine whether UNC-45A co-localizes with MTs in interphase and whether its localization pattern resembles the periodic pattern seen *in vitro* and *in vivo*. For these localization and co-localization studies, we used the same Proteintech anti-UNC-45A antibody. This antibody we also we previously show to be specific in IF, by performing IF staining in shRNA scramble versus shRNA knockdown metaphase cells (Supplementary Figure 3[5]) stained with anti-UNC-45A antibody only (no alpha-tubulin staining).

The first set of experiments was done with RFL-6 cells because they have a very distinct MT network that facilitates analysis of MT and MT associated proteins. Here, cells were stained with either anti-alpha-tubulin alone followed by secondary anti -Alexa Fluor 594 (Figure 4A, *left panel*) or anti-UNC-45A alone followed by secondary anti-FITC (Figure 4A, *right panel*). We found that the pattern of UNC-45A subcellular localization was similar to the one of tubulin suggesting that UNC-45A could be a MT localizing protein in interphase. Importantly, secondary anti-FITC only or secondary anti -Alexa Fluor 594 only had little to no fluorescence when images where taken with the same exposure time indicating that the staining is not due to secondary antibody alone (Figure 4B).

**Figure 4.**
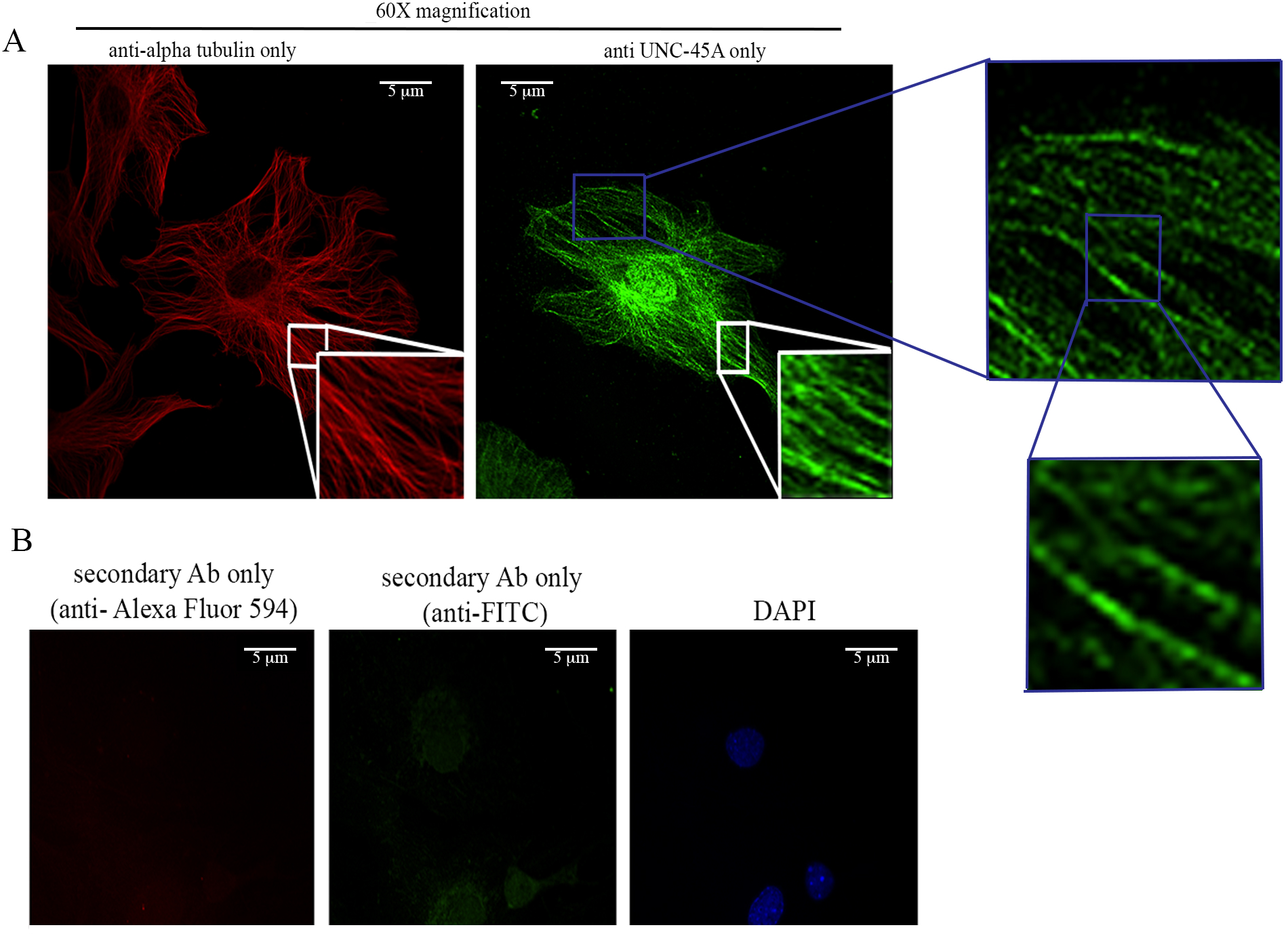
UNC-45A has a localization pattern similar to the one of alpha tubulin in interphase cells. A. Representative images of tubulin (red) and of UNC-45A (green) in RFL-6 cells. B. Representative images of DAPI, and secondary anti-FITC or anti-Alexa Fluor 594 only. All images were taken using the same exposure time. Cells were fixed and permeabilized with methanol.

Next, we used confocal microscopy to determine whether endogenous UNC-45A co-localizes with MTs in RFL-6 cells stained for anti-UNC-45A and anti-alpha-tubulin following their relative conjugated antibodies. As shown in Figure 5A, we observed a clear co-localization of UNC-45A with MTs. Importantly, we confirmed that the red laser does not emit any green signal when the green laser is off and that the green laser does not emit any red signal when the red laser is off (Figure 4B). This, in addition to the fact that FITC and Alexa Fluor 594 are far apart in the excitation fluorescence spectra and that images were taken sequentially, indicates that UNC-45A co-localizes with MTs.

**Figure 5.**
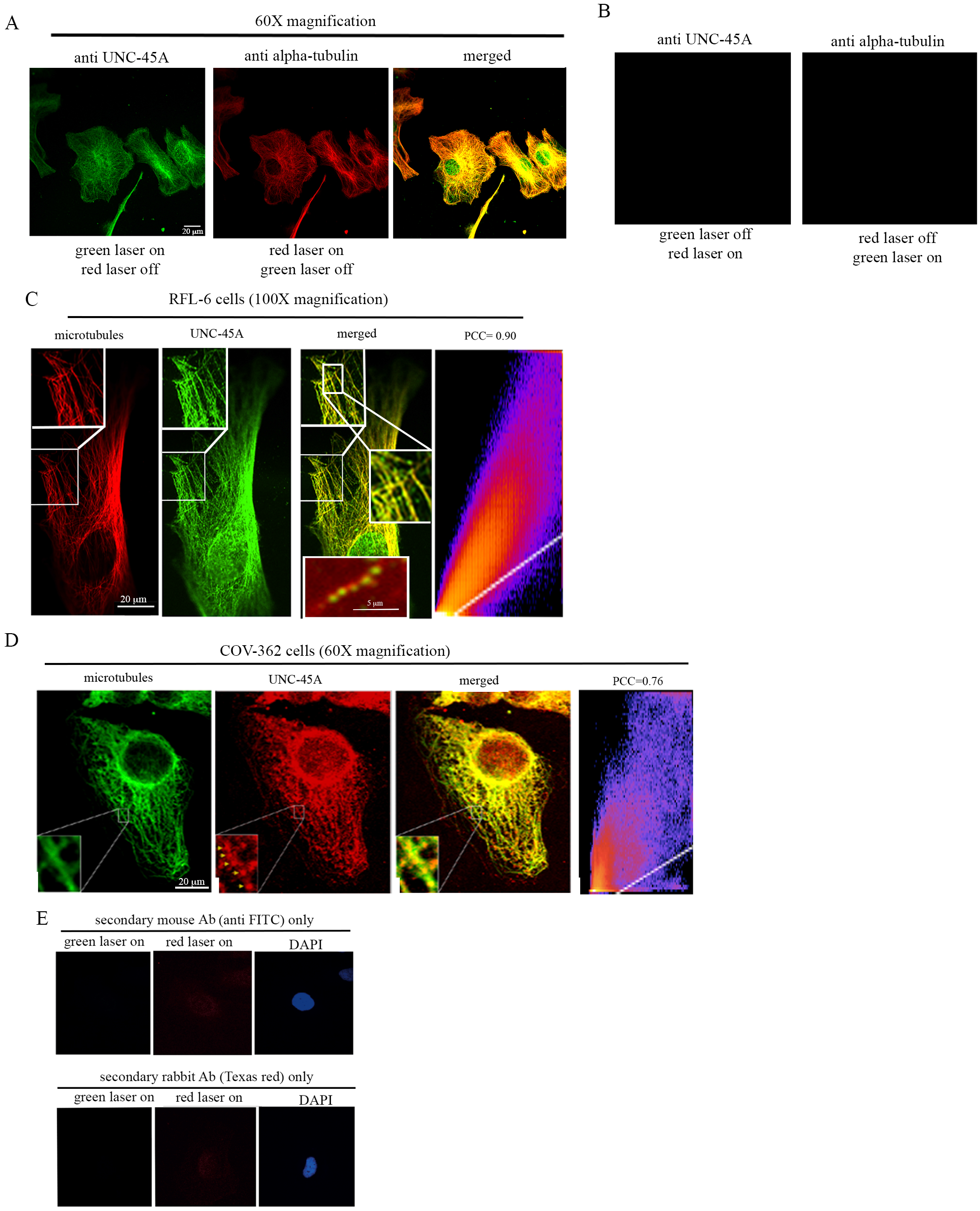
UNC-45A has a punctated localization pattern on interphase MTs *in vivo*. A. Representative images of UNC-45A (green) and tubulin (red) of in RFL-6 cells. Yellow is the merged image. B. Representative images of “red laser on” that does not emit any green signal when the green laser is off and that the “green laser on” does not emit any red signal when the red laser is off. Images where taken using the same exposure time. Images were taken with 60X lens. C. Representative images of UNC-45A (green) and tubulin (red) in interphase RFL-6 cells. Yellow is the merged image. Inset in the merged image is example image of paclitaxel-stabilized MT (red) and UNC-45A-GFP (green). Images were taken with 100X lens. Pearson’s correlation coefficient (PCC) of 0.90. D. Representative images of tubulin (green) and UNC-45A (red) in COV-362 interphase cells. Images were taken with 60X lens. Pearson’s correlation coefficient (PCC) of 0.76. E. Representative images of DAPI, and secondary anti-FITC or Texas-red antibodies only. For these co-localization studies cells were fixed and permeabilized with methanol which is the fixative of choice to for visualizing MTs and their associated proteins. Other fixation methods including formaldehyde and glutaraldehyde and are known to not preserve MTs well and/or interfere with antigen binding (from “*Fluorescence Procedures for the Actin and Tubulin Cytoskeleton in Fixed Cells*”-Louise Cramer and Arshad Desai and “*Fluorescence Microscopy of Microtubules in Cultured Cells*”-Microtubules Protocols Methods in Molecular Medicine).

To gain better visualization of the pattern of UNC-45A localization on MTs, we performed the same experiment and took images using a 100X lense. As shown in Figure 5C, UNC-45A co-localizes with MTs *in vivo*, with a puntacted pattern similar to the one we have previously described *in vitro* on purified MTs [5] (inset in the merged image) and with a Pearson’s correlation coefficient (PCC) of 0.90. The same co-localization of UNC-45A and MTs with a punctated pattern was investigated in interphase COV-362 cells. For these experiments, we used the same primary antibodies as for RFL-6 cells (anti-UNC-45A, rabbit and anti-alpha-tubulin, mouse) and swapped the fluorophores in the secondary antibodies. As show in Figure 5D we observed a similar co-localization and pattern between UNC-45A and MTs in COV-362 cells with a Pearson’s correlation coefficient (PCC) of 0.76. As shown in Figure 5E, secondary antibodies alone had minimal background.

## Discussion

Most of the studies published so far on the mammalian isoform UNC-45A have focused on its role as direct or indirect regulator of actomyosin contractility. This includes studies from our laboratories showing that UNC-45A co-localizes with NMII in mammalian cells including cancer cells, NK cells and neurons [17, 19, 20]. This also includes a study from Dr. Lappalainen’s group showing that the UCS domain of UNC-45A co-localizes with stress fibers in the U2OS cell line where it promotes myosin folding and stress fibers assembly [6].

More recently we and others have shown that UNC-45A has independent functions in actomyosin and MT systems. This includes work published by Dr. Chadli’s group showing that in addition to being a cytoplasmic protein, UNC-45A is also present in the nuclei of cancer cells where it regulates the transcription of the mitotic kinase NEK7 [21]. This also includes: *a)* work from the same Chadli’s group showing that UNC-45A, co-localizes and biochemically co-fractionates with gamma tubulin and regulates centrosomal positioning [15], *b)* work from Borner’s group on quantitative subcellular proteomic analysis of HeLa cells showing that UNC-45A is in the subcellular fractions where other well know MT-associated and destabilizing proteins are found, including katanin and MCAK [7], and *c)* work from our group showing that UNC-45A is a MAP with MT-destabilizing activity that binds and destabilizes MT in absence of myosin II [5].

Here we initially validated two of the most commonly used anti-UNC-45A antibodies and show that while they are both remarkably clean, the rabbit polyclonal anti-UNC-45A from Proteintech seems to be more specific as compared to the mouse polyclonal anti-UNC-45A from Abnova we tested. This is the same rabbit antibody we had previously used for IHC and IF studies to show that UNC-45A is a mitotic spindle-associated protein [5]. We and other had previously shown that UNC-45A is expressed differentially in cancer versus normal cells and that its upregulation at both protein and RNA levels correlates with the severity of the disease in ovarian, breast and melanoma patients [14, 17] and the Cancer Genome Atlas. Subcellular fractions studies conducted in HeLa cells shows that UNC-45A is very abundant at 400nM [7]. Here we show that of all the cell lines we tested, HeLa was indeed the one with the highest levels followed by the ovarian cancer cell lines OVTOKO, RMGI and ES-2. Interestingly, the OVTOKO ovarian cancer cell line is highly resistant to paclitaxel and can be sensitized to the drug by increasing MT stability via inhibition of the MT stabilizing protein SCLIP [22]. The potential therapeutic applications of this discovery include consideration of strategies such as sequential or concurrent use of such MT stabilizing protein inhibitors with other drugs in patients with paclitaxel-resistant ovarian cancers. This finding provides opportunity for further studies to investigate this potential novel avenue.

Consistent with the subcellular fractions studies [7] we had previously shown that UNC-45A co-fractionated with MTs in the ovarian cancer cell line COV-362. Here we confirmed and expanded our previous studies and showed that the increase in UNC-45A in the pellet fraction following taxol-treatment is due to its binding to MTs and not to actin or myosin II. In this scenario, binding of UNC-45A to both the actomyosin system and MTs is not mutually exclusive.

As far as UNC-45A subcellular localization, consistent with our previous studies conducted in synchronized cell line UNC-45A was found to localize with mitotic spindle in clinical specimens ovarian cancer, was predominantly expressed in cancer cells versus stromal cells and had predominant cytoplasmic localization *in situ* in cancer cells. Interestingly, while we and others have previously shown that UNC-45A is mostly a cytoplasmic, perinuclear protein a new study from Dr. Chadli’s laboratory shows that in clinical specimens, part of UNC-45A is found in the nucleus where it regulates NEK7 activity [21]. Consistent with our previous studies of UNC-45A co-localization with MTs in metaphase cells [5], here we show that UNC-45A co-localizes with interphase MTs in a variety of cells including HeLa, RFL-6 and COV-362.

The fact that we and other have previously “missed” the localization of UNC-45A with MTs is not surprising for a number of reasons. First, not all cells have a MT network suitable for MT localization and co-localization studies, in fact, most mammalian cells do not. This is particularly true if, like in the case of UNC-45A [5], the MAP of interest binds the MT lattice rather than to the MT plus ends [23]. Second, while mitotic spindles are formed by bundles of MT and therefore are relatively easy to detect, the percentage of mitotic and metaphase cells in an unsynchronized population in generally less than 5% at any given time *in vitro* and *in vivo* [24]. Interphase MT on the other hand, are individual and more difficult to detect and do require confocal microscopy for co-localization studies. Third, fixation methods other than cold methanol (which we have used here and previously to show that UNC-45A is a MAP[5]) may not be suitable for MT localization or co-localization studies because their either do not preserve MTs well, or interfere with antigen binding. With regards to this point, our work on co-localization of UNC-45A with NMII in NK cells and neurons [17, 19, 20] as well work from Drs. Lappalainen and Chadli’s groups were all done using formaldehyde fixation because that is the preferred method for studying the actomyosin system. Additionally, while our previous work on UNC-45A co-localization with myosin II in cancer cells was done using methanol as a fixation method, oflocalization with MT was not evaluated and confocal microscopy was not used. Last but not least, no more than 61 papers to date are published on this highly conserved protein, clearly indicating that we are just now uncovering its multiple functions in cells.

The fact that a protein can have a dual role in regulating actomyosin and MT stability is not mutually exclusive and well documented in scientific literature. For instance, one of the most well-known cell cycle regulators, the cyclin-dependent kinase inhibitor 1B (p27), is both a MT- and a NMII-associated and co-localizing protein and has been show to regulate both NMII activity and MTs stability in normal and cancer cells[25–31]. If fact, while the interactions between actomyosin and MT systems are well known, the effects of actomyosin system on MT are generally modest and in reverse direction given that of myosin II leads to modest increase in MT stability[32].

Additionally, many MTs destabilizing proteins including katanin localize at the centrosomes, at mitotic spindles, and at the actomyosin contractile ring where they participate in spatiotemporal regulation of cytokinesis[33] [34].

Lastly, UNC-45A abundance, along with the duality of its functions within a cell as both an MT- and actinomysoin-associated protein, suggest that the activity of UNC-45A must be tightly regulated. Two large phosphoproteomic screens identified a phosphothreonine residue at position 15 in UNC-45A [35, 36]. Given that phosphorylation is a well-documented mechanism to regulate the activity of microtubule-associated proteins [37–39] it is likely that a complex system of kinases and phosphatases cooperate to fine-tune the activity of UNC-45A in cells.

Taken together we believe that understanding the multiple roles of UNC-45A in regulating cytoskeletal dynamics is key for broadening our understanding of cell biology in both the context of health and disease.

## Materials and Methods

### Cell culture

ES-2, TOV-21G, SKOV-3 and HeLa cells were purchased from the American Type Culture Collection (ATCC) and cultured in DMEM (Thermo Fisher) supplemented with 10% FBS as previously described [40]. RFL-6 cell were purchased from the American Type Culture Collection (ATCC) Ham’s F-12K medium (Thermo Fisher) supplemented with 20% FBS. The ovarian cancer cell line COV-362 was a generous gift from Dr. Panagiotis A. Kostantinopoulos (Dana-Farber Cancer Institute, Boston, MA) and was cultured in DMEM (Thermo Fisher) supplemented with 10% FBS. The JHOC5 and OVISE cells were a generous gift from Dr. Tian-Li A. Wang (Johns Hopkins Medical School, Baltimore, MD) and were cultured in RPMI supplemented with 10% FBS. The OVTOKO ovarian cancer cell line was a generous gift of Dr. David Huntsman at the University of British Columbia (Vancouver, Canada) and was cultured in DMEM (Thermo Fisher) supplemented with 10% FBS. Cells routinely tested negative for mycoplasma.

### Human subjects

Archival tissues were used with the Institutional Review Board approval.

### Immunohistochemistry

Five-micron thick formalin-fixed, paraffin-embedded sections were deparaffinized and rehydrated by sequential washing with xylene, 100% ethanol, 95% ethanol, 80% ethanol, and PBS. For antigen retrieval, slides were immersed in Reveal Decloaker (Biocare Medical, Concord, CA) and steamed for 30 min at 100 degrees C. Endogenous peroxidase activity was blocked with 3% H_2_O_2_ for 10 min. After washing with PBS, slides were blocked with 10% normal goat serum in PBS for 10 min at room temperature, followed by incubation with rabbit anti-human polyclonal UNC-45A antibody (Proteintech Group Inc) at a concentration of 1:200 in blocking solution overnight at 4 degrees C. After washing twice with PBS, slides were incubated with a biotinylated anti-rabbit secondary antibody conjugated (10 min) and streptavidin/horseradish peroxidase (10 min; Dako), followed by 3,3-diaminobenzidine (Phoenix Biotechnologies) substrate for 3 min. Slides were lightly counterstained with Gill No. 3 hematoxylin (Sigma) for 60 s, dehydrated, and coverslipped.

### Single cell RNA sequencing

Patients were enrolled in the study upon diagnosis of ovarian cancer. The study was approved by the University of Minnesota’s IRB and all patients provided informed consent. Fresh tumor samples were digested with an enzyme cocktail following standard protocols to produce a single cell suspension (Miltenyi GentleMacs Dissociator Kit, Auburn, CA). Single cells were processed for sequencing using Gel Emulsion Beads using the 10X Genomics Chromium Single Cell 3’ kit (10X Genomics, Pleasanton, CA). Sequencing was performed on an Illumina HiSeq 2500. Raw sequence processing, de-multiplexing and mapping was performed using CellRanger software (10X Genomics, Pleasanton, CA). Quantification, cell type assignment and TSNE plot visualization was done using the Seurat R package (Seurat v3.0.0.9000. Satija, et al., Nature Biotechnology 2015)

### Lentiviral-mediated UNC-45A knockdown in cells

For UNC-45A silencing, scramble and UNC-45A shRNAs lentiviral supernatant were prepared and used to infect HeLa, as we have previously described [5, 20].

### Antibodies and chemicals

Rabbit anti-UNC-45A raised against the C-terminus of the human UNC-45A (Protein Tech 1956-1-AP) was used for Western blot and immunofluorescence at the concentration recommended by the manufacturer (1:1000 dilution for Western blot and 1:200 dilution for immunofluorescence) and previously used in our laboratory [5]. Mouse polyclonal anti UNC-45A raised against full length UNC-45A (Abnova) was used for Western blot at the at the concentration recommended by the manufacturer (1:300 dilution) and as previously described[6]. Anti-NMII (LC) (Sigma), anti-actin (Sigma), and anti-alpha-tubulin (Abcam) were used at the concentration recommended by the manufacturers. Peroxidase-linked anti-mouse immunoglobulin G and peroxidase-linked anti-rabbit immunoglobulin G (both GE Healthcare Bio-Sciences, Pittsburgh, PA). Alexa Fluor 594-conjugated Donkey Anti-Mouse IgG (1:250) and FITC-conjugated Goat Anti-Rabbit IgG (1:200; both Jackson ImmunoResearch Laboratories, West Grove, PA). Texas-Red conjugated Goat Anti-Mouse IgG (1:250; both Jackson ImmunoResearch Laboratories, West Grove, PA), and FITC-conjugated Goat Anti-Mouse IgG (1:250; both Jackson ImmunoResearch Laboratories, West Grove, PA).

### Western blot analysis

Total cellular protein (7–50 µg) from each sample was separated by SDS-PAGE, transferred to PVDF membranes and subjected to Western blot analysis using the specified antibodies. Amido black staining was performed to confirm equal protein loading.

### Recombinant protein

GFP-UNC-45A and its mutants, were cloned into pGEX-2TK to generate the GST-GFP-UNC-45A protein. The protein was expressed in Rosetta (DE3) pLysS and following GST removal it was affinity purified and dialyzed as we have previously described[5].

### Microtubule cosedimentation assay in cells

Cells were lysed using 1% NP-40 lysis buffer containing 150 mM NaCl, 50 mM Tris-HCl pH 7.8, and protease inhibitor cocktail. The lysates were treated with either control DMSO or 1μM of taxol and incubated at 37°C for an hour. Lysates were spun at 15,000rpm for 30 minutes at room temperature and the supernatant and pellet fractions were separated by SDS-PAGE and analyzed via Western blotting. Extent of UNC-45A, alpha-tubulin, actin and myosin II in the different cell compartment (supernatant or pellet) with or without taxol treatment was determined by densitometric analysis using imageJ software.

### Immunofluorescence microscopy, image acquisition, and analysis

For localization and co-localization analysis of UNC-45A and MTs cells were fixed in cold methanol for 5 minutes at −20°C. After blocking with 5% BSA in PBST, cells were stained with anti-UNC-45A (rabbit) and anti-α-tubulin (mouse) primary antibodies followed by FITC- (rabbit) or Alexa Fluor 594- (mouse), or FITC- (mouse) or Texas red- (rabbit) conjugated secondary antibodies and analyzed via confocal fluorescence microscopy. Images were taken with an Olympus BX2 upright microscope equipped with a Fluoview 1000 confocal scan head. A UPlanApo N 100X/1.42 NA objective was used. FITC was excited with a 488 nm laser and emission collected between 505 and 525 nm. For Alexa Fluor a 594 nm laser was used for excitation and emission collected between 560 and 660 nm. Images were taken with sequential excitation. Co-localization analysis was done using the Fiji software Coloc 2 plugin and Pearson’s correlation coefficient (PCC) and calculated as previously described [41].

Images were obtained on an Axiovert 200 microscope (Zeiss, Thornwood, NY) equipped with a high-resolution CCD camera. All images were obtained using identical camera, microscope, and imaging criteria such as gain, brightness, contrast, and exposure time.

### Statistical analysis

Results are reported as mean ± Standard Deviation of three or more independent experiments. Unless otherwise indicated, statistical significance of difference was assessed by two-tailed Student’s t using Prism (V.4 Graphpad, San Diego, CA) and Excel. The level of significance was set at p<0.05.

## Acknowledgments

We thank Guillermo Marques (University of Minnesota Imaging Center) for assistance with image analysis. This work was supported by Department of Defense Ovarian Cancer Research Program Grant OC160377, the Minnesota Ovarian Cancer Alliance and the Randy Shaver Cancer Research and Community Fund grant to Martina Bazzaro, by the Ovarian Cancer Research Fund Grant to Boris Winterhoff and by the University of Minnesota Grand Challenges Grant to Tim Starr and Boris Winterhoff. Emil Lou is supported by the Litman Family Fund for Cancer Research, Randy Shaver Cancer Research and Community Fund, Minnesota Masonic Charities, the Masonic Cancer Center (Grant # P30 CA77598) and Department of Medicine, Division of Hematology, Oncology and Transplantation, University of Minnesota to Emil Lou. The funders had no role in study design, data collection and analysis, decision to publish or preparation of the manuscript. The authors declare no conflict of interests.

